# Moxidectin and ivermectin inhibit SARS-CoV-2 replication in Vero E6 cells but not in human primary airway epithelium cells

**DOI:** 10.1101/2021.05.17.444467

**Authors:** Nilima Dinesh Kumar, Bram M. ter Ellen, Ellen M. Bouma, Berit Troost, Denise P. I. van de Pol, Heidi H. van der Ende-Metselaar, Djoke van Gosliga, Leonie Apperloo, Orestes A. Carpaij, Maarten van den Berge, Martijn C. Nawijn, Ymkje Stienstra, Izabela A Rodenhuis-Zybert, Jolanda M. Smit

## Abstract

Antiviral therapies are urgently needed to treat and limit the development of severe COVID-19 disease. Ivermectin, a broad-spectrum anti-parasitic agent, has been shown to have anti-SARS-CoV-2 activity in Vero cells at a concentration of 5 µM. These *in vitro* results triggered the investigation of ivermectin as a treatment option to alleviate COVID-19 disease. In April 2021, the World Health Organization stated, however, the following: “the current evidence on the use of ivermectin to treat COVID-19 patients is inconclusive”. It is speculated that the *in vivo* concentration of ivermectin is too low to exert a strong antiviral effect. Here, we performed a head-to head comparison of the antiviral activity of ivermectin and a structurally related, but metabolically more stable, moxidectin in multiple *in vitro* models of SARS-CoV-2 infection, including physiologically relevant human respiratory epithelial cells. Both moxidectin and ivermectin exhibited antiviral activity in Vero E6 cells. Subsequent experiments revealed that the compounds predominantly act on a step after virus cell entry. Surprisingly, however, in human airway-derived cell models, moxidectin and ivermectin failed to inhibit SARS-CoV-2 infection, even at a concentration of 10 µM. These disappointing results calls for a word of caution in the interpretation of anti-SARS-CoV-2 activity of drugs solely based on Vero cells. Altogether, these findings suggest that, even by using a high-dose regimen of ivermectin or switching to another drug in the same class are unlikely to be useful for treatment against SARS-CoV-2 in humans.

## 1. Introduction

Within less than 1.5 year, the pandemic SARS coronavirus 2 (SARS-CoV-2) has infected over 153 million individuals and resulted in over 3.2 million deaths worldwide (Sohrabi et al., 2020; WHO, 2021a; Zhou et al., 2020). The social and economic burden of this still ongoing pandemic is staggering and besides vaccine development, it is of utmost importance to develop therapeutic interventions to reduce disease symptoms. To date, multiple compounds have been shown to exert SARS-CoV-2 antiviral activity *in vitro* and several compounds have reached clinical trials (Chan et al., 2020; Zhang et al., 2020). Remdesivir and hydroxychloroquine were thought to be effective early in the pandemic but after a careful evaluation in an interim solidarity trial, the WHO released a conditional yet strong recommendation against the usage of these drugs as no impact on overall mortality was observed (Consortium et al., 2021). Corticosteroids is currently (April 29^th^, 2021) the only therapeutic agent strongly recommended by the WHO for the treatment of severe and critical COVID-19 patients (Siemieniuk et al., 2020). These guidelines, however, differ from National Institute of Health (NIH) guidelines which recommend anti-SARS-CoV-2 monoclonal antibodies for selective patients with mild to moderate disease and remdesivir, dexamethasone and tocilizumab either individually or in combination based on the disease severity (NIH, 2021).

Ivermectin, a macrocyclic lactone of the avermectin subfamily, is a broad-spectrum anti-parasitic agent approved by the United States Food and Drug Administration (FDA) and European Medicines Agency (EMA) for prophylactic and therapeutic usage in some animal species and for selective treatments in humans (EMA, 2009, 2017; Makhani et al., 2019; Prichard et al., 2012). In recent years, ivermectin has also been shown to have antiviral activity *in vitro* towards several viruses including Zika virus (ZIKV) and dengue virus (DENV) (Gotz et al., 2016; Lundberg et al., 2013; Tay et al., 2013; Varghese et al., 2016; Wagstaff et al., 2012). In addition, Caly and colleagues (Caly et al., 2020) showed that ivermectin effectively inhibits SARS-CoV-2 infection in Vero/hSALM cells. The success of ivermectin as an antiviral agent *in vitro* has, however, been proven difficult to translate to *in vivo* settings. For example, ivermectin failed to protect against lethal ZIKV challenge in mice (Ketkar et al., 2019) and did not reduce DENV viremia in phase III clinical trials (Yamasmith et al., 2018). It is speculated that the *in vivo* concentration of ivermectin is too low to exert its antiviral effect (Bray et al., 2020; Momekov and Momekova, 2020). Notably, despite the known limitation in achieving high ivermectin concentrations in humans (Pena-Silva et al., 2020; Schmith et al., 2020) and the limited *in vitro* proof of anti-SARS-CoV-2 activity (Caly et al., 2020), 65 ivermectin clinical trials (April 29^th^, 2021) are registered as treatment intervention against COVID-19 (clinicaltrials.gov, 2021). Recently, upon review of the latest trail results, many regulatory authorities including WHO issued a recommendation stating “to not use ivermectin in patients with COVID-19 except in the context of a clinical trial” as the available evidence to support its usage is uncertain (Chaccour et al., 2021; Gonzalez et al., 2021; Lopez-Medina et al., 2021; WHO, 2021b).

Moxidectin, a macrocyclic lactone belonging to the milbemycin subfamily and structurally related to ivermectin, is a broad-spectrum anti-parasitic used in veterinary medicine. In addition, it has recently been approved for human use for the prevention of river blindness, a disease caused by the parasite *Onchocerca volvulus* (Cobb and Boeckh, 2009; Milton et al., 2020; Prichard and Geary, 2019). Importantly, moxidectin has been shown to have superior drug disposition properties to ivermectin such as a longer half-life and higher efficacy in animals and humans (Opoku et al., 2018; Prichard and Geary, 2019). To date, antiviral potential of moxidectin towards SARS-CoV-2 has not been evaluated and no clinical trials are registered for moxidectin as COVID-19 treatment.

In this study, we evaluated the antiviral activity of moxidectin in direct comparison with ivermectin towards SARS-CoV-2 infection *in vitro*. We utilized the commonly used African green monkey kidney (Vero E6) cells to compare the effective antiviral concentrations and to evaluate the longevity and mechanistic properties of the compounds. To verify the results in a physiological more relevant model system we subsequently tested the antiviral activity of both drugs in the human lung epithelial cells (Calu-3) and in primary human bronchial epithelial cells (PBECs) grown under air-liquid interface (ALI) culture conditions (Leist et al., 2020).

## 2. Materials and Methods

### 2.1. Chemicals and reagents

Ivermectin (Sigma-Aldrich, MO, USA) and moxidectin (European pharmacopeia reference standard, Strasbourg, France) were dissolved in absolute ethanol (EtOH) (Sigma-Aldrich, MO, USA) to a final concentration of 5 mM and stored at -20 °C. The maximum final EtOH concentration corresponded to 0.2% in all experiments.

### 2.2. Virus stock and titration

SARS-CoV-2 strain /NL/2020 (European Virus archive global (EVAg), 010V-03903) was produced in Vero E6 cells. The cells were infected at a multiplicity of infection (MOI) of 1 and 48 hours post-infection (hpi) supernatants containing progeny virions were harvested, centrifuged, aliquoted and stored at -80 °C. The obtained virus was passaged twice prior to usage for experiments. The infectious virus titer was determined by a plaque assay on Vero E6 cells. Briefly, Vero E6 cells were infected for 2 h with 10-fold serial dilutions of samples following, which, cells were overlaid with 1:1 mixture of 2% agarose (Lonza, Basel, Switzerland) and 2X MEM medium. At 72 hpi, the plaques were fixed using 10% formaldehyde (Alfa Aesar, Kandel, Germany) and stained using crystal violet (Sigma-Aldrich, MO, USA). Infectious titers are stated as plaque forming units (PFU) per ml. One plaque in 10-fold dilution corresponds to 150 PFU/ml and was set as the threshold of detection for all experiments.

### 2.3. Cell culture

The African green monkey Vero E6 cell line (ATCC CRL-1586), kindly provided by Gorben Pijlman (Wageningen University, Wageningen, the Netherlands) was maintained in Dulbecco’s minimal essential medium (DMEM) (Gibco, Paiskey, UK), high glucose supplemented with 10% fetal bovine serum (FBS) (Lonza, Basel, Switzerland), penicillin (100 U/mL), and streptomycin (100 U/mL) (Gibco, Paiskey, UK). The human lung epithelial cell line Calu-3 (ATCC HTB-55) was maintained in DMEM F-12 (Lonza, Basel, Switzerland) supplemented with 10% FBS, 1% Glutamax (Thermo Fisher Scientific, Inc., Waltham, MA, USA), 1% non-essential amino acid (Thermo Fisher Scientific, Inc., Waltham, MA, USA), penicillin (100 U/mL), and streptomycin (100 U/mL). All cells were mycoplasma negative and maintained at 37 °C under 5% CO_2_. Primary human bronchial epithelial cells (PBECs) were cultured from bronchial brushing obtained by fibreoptic bronchoscopy performed using a standardized protocol during conscious sedation (Heijink et al., 2007; Vieira Braga et al., 2019). The medical ethics committee of the University Medical Center Groningen approved the study (METC 2019/338), and all subjects gave their written informed consent. The donors were 3 male and 2 female non-smoking healthy control volunteers, with a normal lung function (FEV/FVC > 70%, FEV_1_ > 90% predicted) and absence of bronchial hyperresponsiveness to methacholine (PC_20_ methacholine > 8 mg/ml). PBECs were cultured and fully differentiated under ALI conditions in transwell inserts, as previously described (Heijink et al., 2010).

### 2.4. Cytotoxicity assays

#### 2.4.1. MTS assay

MTS assay, to determine cytotoxicity, was performed using the CellTiter 96^®^ AQueous One Solution Cell Proliferation Assay kit using manufacturer’s instructions from Promega (Madison, WI, USA). Vero E6 cells were seeded in 96-well plates at a density of 10,000 cells/well. Following day, cells were treated with increasing concentrations of moxidectin or ivermectin ranging from 0 to 80 µM for 8 h or with 10 µM of both compounds for 60 h along with an equivalent volume of EtOH. Thereafter, 20 µl of the MTS/PMS reagent was added to each well and the cells were further incubated at 37 °C for 2 h. Subsequently, 10% SDS solution was added to stop the reaction and the absorbance was measured at 490 nm using a microplate reader (BioTek, Winnooski, VT). Calu-3 cells were seeded in 96-well plate at a density of 40,000 cells/well. Cells were treated with 5 and 10 µM moxidectin, ivermectin or an equivalent volume of EtOH for 8 h. Same steps were followed as described above for Vero E6 cells. Cytotoxicity was determined based on the following formula:

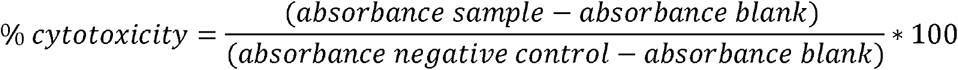

#### 2.4.2. LDH cytotoxicity assay

LDH assay was performed using the CyQUANT™ LDH Cytotoxicity Assay Kit (Thermo Fisher Scientific, Inc., Waltham, MA, USA). PBECs cultured under ALI conditions were treated with 10 µM moxidectin, ivermectin or an equivalent volume of EtOH at the basolateral side for 48 h at 37 °C. Thereafter, LDH release was determined at the apical side. Hereto, 200 µL of OptiMEM (Gibco, Paiskey, UK) was added to the apical side 30 min prior to the harvest. The apical harvest was clarified by centrifuging at 2000 x g at 4 °C. Levels of LDH were determined according to manufacturer’s instructions in all experimental conditions. The absorbance was measured at 490 and 680 nm using a microplate reader (BioTek, Winnooski, VT). Absorbance at 680 nm was subtracted from the absorbance at 490 nm and cytotoxicity was calculated as described below:

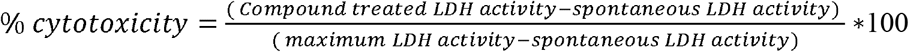

#### 2.4.3. Live/dead staining and flow cytometry

PBECs cultured under ALI conditions were treated with 10 µM moxidectin, ivermectin or an equivalent volume of EtOH at the basolateral side for 48 h at 37 °C. Subsequently, cells were harvested by trypsinization and stained with fixable viability dye eFluor780 (Thermo Fisher Scientific) for 20 min at 4 °C. Next, cells were washed with FACS buffer (1X PBS, 2% FBS, 1% EDTA), centrifuged and fixed with 4% PFA for 10 min at 4 °C. After fixation, cells were washed, centrifuged and resuspended in FACS buffer. Flow cytometry analyses was performed using the LSR-2 flow cytometer (BD Biosciences, San Jose, CA, USA) and data was further analyzed using Kaluza analysis software, version 2.1 (Beckman Coulter, Fullerton, CA, USA).

### 2.5. Antiviral assay in Vero E6 and Calu-3

Vero E6 cells were seeded at a density of 1.3 ×10^5^ cells/well in 12-well plates. Next day, the medium was replaced with 0.25 mL of DMEM (2% FBS) medium containing the virus inoculum (MOI 1) and in presence of increasing concentration of compounds or the equivalent volume of EtOH. Following 2 h adsorption at 37 °C, the virus inoculum was removed, after which the cells were washed twice and fresh DMEM media (10% FBS) containing the compounds or EtOH was added. At 8 hpi, cell supernatants were harvested and titrated using plaque assay. For the durability assay, Vero E6 cells were infected with SARS-CoV-2 at MOI 0.01 and treated with 10 µM of moxidectin, ivermectin or the equivalent volume of EtOH. Supernatants were collected at 16, 24, 40 and 60 hpi. A lower MOI was used to allow multiple rounds of infection. Samples were titrated using plaque assay. Calu-3 cells were seeded at a density of 2 ×10^5^ cells/well in 24-well plates. At 48 h post-seeding, infection was performed in 0.2 mL of DMEM (2% FBS) medium containing virus inoculum (MOI 1) and 5 or 10 µM of moxidectin, ivermectin or the corresponding volume of EtOH. Cell supernatants were harvested at 8 hpi and titrated using plaque assay.

### 2.6. Antiviral assay in primary bronchial epithelial cells

Following 3 weeks of ALI culture, human primary bronchial epithelial cells (PBECs) were washed twice with OptiMEM to remove excess mucus. At the time of infection, 10 µM moxidectin, ivermectin or the equivalent volume of EtOH was added at the basolateral side of the insert (12-well). The apical side was inoculated with SARS-CoV-2 at MOI 5. At 2 hpi, cells were washed twice with OptiMEM and left on air at 37 °C until collection. Thirty minutes prior to collection (12, 24 and 48 hpi), 150 µL of OptiMEM was added to the apical side of the ALI cultures and virus was harvested by incubating for 30 min at 37 °C. Infectious virus titers were determined using plaque assay.

### 2.7. Virucidal assay

SARS-CoV-2 particles (2.5 ×10^5^ PFUs) were incubated in the presence or absence of 10 µM moxidectin, ivermectin or the equivalent volume of EtOH for 2 h at 37 °C in 300 µl of DMEM media (2% FBS). Infectious virus titers were determined using plaque assay.

### 2.8. Time-of-drug-addition assay

For the time-of-drug-addition experiments, Vero E6 cells were treated with 10 µM moxidectin, ivermectin or the corresponding volume of EtOH either pre, during, or post-inoculation conditions **(Fig. 2B)**. For pre-treatment, cells were incubated with the compounds for 2 h prior to infection. At the time of infection, cells were washed three times and infected with SARS-CoV-2 at MOI 1 for 2 h. At 2 hpi, cells were washed three times with plain DMEM, and media was replaced with DMEM 10% FBS, and incubation was continued until collection. For the “during” condition, the compounds were present together with the virus inoculum, thus only during the 2□h infection time. For the post-inoculation conditions, the compounds were added to the cell culture medium at 2, 4, and 6 h post-inoculation. All supernatants were collected at 8 hpi, clarified by centrifugation and used to quantify the infectious particle titers using plaque assay.

### 2.9. Statistical analysis

The concentration at which moxidectin and ivermectin reduced virus particle production by 50 and 90% is referred to as EC50 and EC90, respectively. Dose-response curves were fitted by non-linear regression analysis employing a sigmoidal model. All data were analyzed in GraphPad Prism software (La Jolla, CA, USA). Data are presented as mean ± SEM. Student T test was used to evaluate statistical differences between treated samples and a p value ≤ 0.05 was considered significant with * p ≤ 0.05, ** p ≤ 0.01 and *** p ≤ 0.001 and ns as non-significant.

## 3. Results

### 3.1 Moxidectin and ivermectin inhibit SARS-CoV-2 infection in Vero E6 cells

First, we assessed the antiviral activity of moxidectin towards SARS-CoV-2 in the African green monkey kidney epithelial Vero E6 cell line and compared that to the efficacy of ivermectin. Vero E6 is highly permissive to SARS-CoV-2 infection (Matsuyama et al., 2020) and thus commonly used in studies investigating virus-host interactions and potential antiviral drugs. Prior to assessing the antiviral efficacy, we determined the cellular cytotoxicity of moxidectin and ivermectin in Vero E6 cells. A clear dose-dependent cytotoxicity was observed **(Fig. S1)**. The highest non-toxic dose was set at 10 µM for subsequent experiments. At this concentration, the cell viability was above 90% for both moxidectin **(Fig. S1A)** and ivermectin **(Fig. S1B)**. Next, Vero E6 cells were infected with SARS-CoV-2 at a multiplicity of infection (MOI) of 1 in presence of 10□µM moxidectin and ivermectin or the equivalent volume of EtOH and virus progeny was determined at 8 hpi. This time point corresponds to 1 cycle of replication (Ogando et al., 2020). SARS-CoV-2 infection under non-treated (NT) conditions led to a production of 4.9□±□0.8 ×10^5^ PFU/mL **(Fig. 1A)**. A comparable titer was observed for the EtOH control (4.3□±□0.7 ×10^5^ PFU/mL), indicating that the solvent does not influence infectious virus particle production. In line with previous results (Caly et al., 2020), ivermectin was found to exert significant antiviral activity towards SARS-CoV-2 in Vero E6 cells **(Fig.1)**. In presence of 10□µM ivermectin, infectious virus particle production was reduced to 1.9□±□0.8 ×10^2^ PFU/mL, which corresponds to a reduction of more than 99.9% when compared to the EtOH control **(Fig. 1A)**. In presence of 10 µM of moxidectin, the infectious virus titer was reduced to 4.3□±□2.0 ×10^2^ PFU/mL which also corresponds to a reduction of 99.9% when compared to the EtOH control **(Fig. 1A)**. Next, we performed a dose-response analysis to determine the EC_50_ and EC_90_ values (i.e. a reduction of 50% and 90% in viral titer, respectively). To this end, Vero E6 cells were infected with SARS-CoV-2 in the presence of increasing concentrations of both moxidectin, ivermectin, and the corresponding amount of EtOH. Moxidectin and ivermectin showed a dose-dependent antiviral activity in Vero E6 **(Fig. 1B)**, with an EC_50_ and EC_90_ of 4.5□µM and 7.2□µM for moxidectin and 1.9□µM and 5.8□µM for ivermectin, respectively **(Fig. 1B)**. The observed EC50 for ivermectin is in line with the previously published value of ∼ 2µM (Caly et al., 2020). Thus, both compounds exhibit potent antiviral effects towards SARS-CoV-2, ivermectin being slightly more potent than moxidectin in this experimental set-up.

**Fig. 1.**
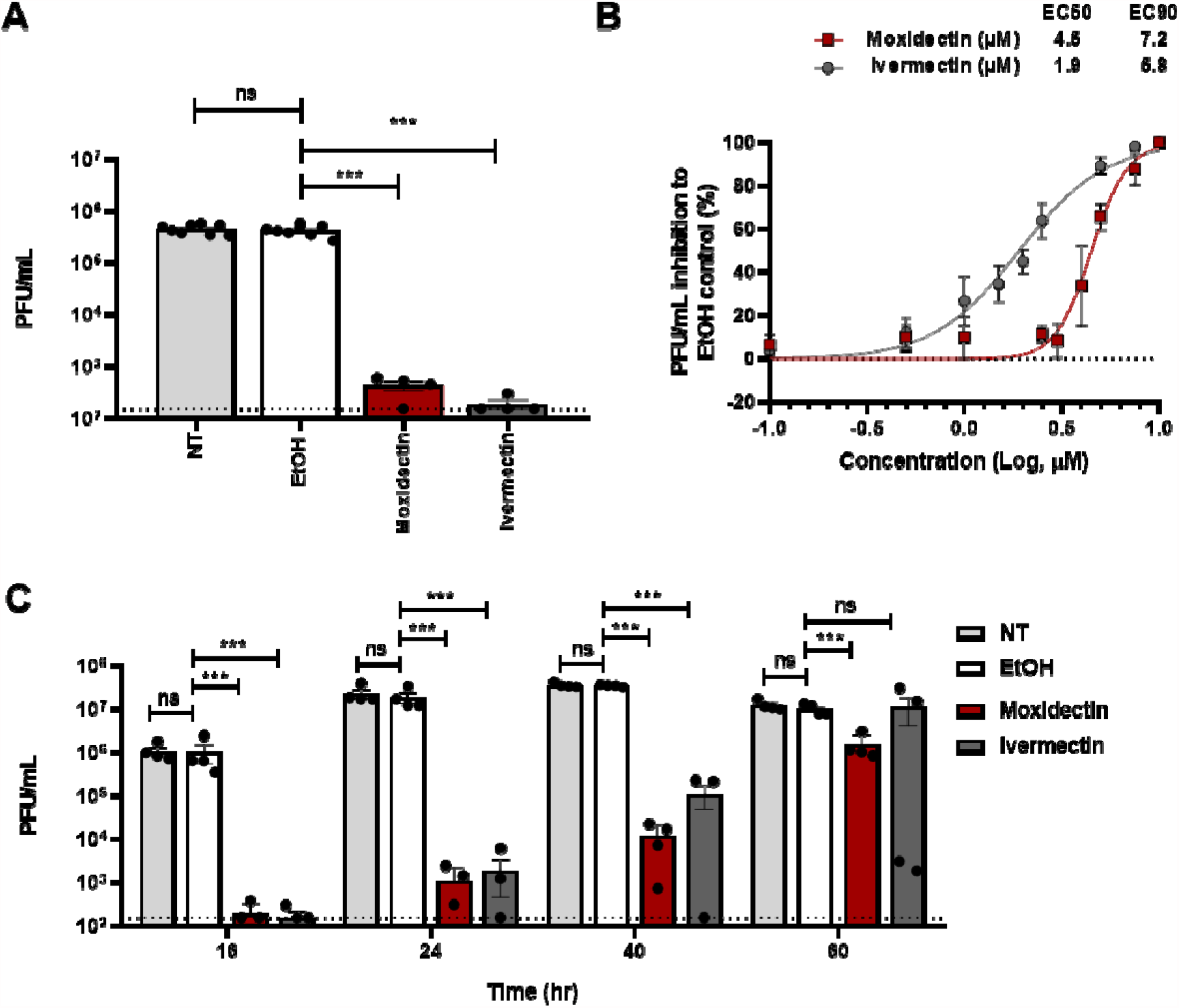
Antiviral activity of moxidectin and ivermectin towards SARS-CoV-2 in Vero E6 cells. Vero E6 cells were treated with 10 μM moxidectin or ivermectin in the presence of SARS-CoV-2 at MOI 1. At 2 h post-infection (hpi), cells were washed twice with plain DMEM medium and new cell culture medium containing the compound was added and incubation was continued for another 6 h. At 8 hpi, cell supernatants were harvested and number of produced infectious virus particles were determined by plaque assay. **(B)** Vero E6 cells were treated with increasing concentrations of moxidectin or ivermectin and infection was continued as described above (A). The EC50 and EC90 values were calculated using GraphPad Prism software. (C) Antiviral effect of 10 µM moxidectin or ivermectin or the corresponding volume of EtOH at 16, 24, 40 and 60 hpi. The dotted lines indicate the detection limit of plaque assay. Each dot represents data from a single independent experiment. Data are represented as mean ± SEM of at least three independent experiments. Statistical analysis was carried out by comparing treated samples with the EtOH control using Student’s t-test *** p < 0.001, ** p< 0.01, * p <0.05 and ns as non-significant.

Given the known increased stability of moxidectin over ivermectin we next evaluated the durability of the antiviral effect for both compounds. To this end, Vero E6 cells were infected with SARS-CoV-2 at MOI 0.01 in presence of 10 µM of moxidectin, ivermectin or corresponding amount of EtOH and the cell supernatants were harvested at 16, 24, 40 and 60 hpi. These time points roughly correspond to 2, 3, 5 and 7 SARS-CoV-2 replication cycles, respectively (Ogando et al., 2020). Importantly, the prolonged incubation of the cells with moxidectin and ivermectin did not influence cell viability as measured with the MTS assay **(Fig. S1C)**. At 40 hpi, 3.5□±□0.4 ×10^7^ PFU/mL were produced at non-treated infection conditions and at this time point virus particle production reached its plateau **(Fig. 1C)**. Comparable titers were observed over time for the EtOH control (3.4□±□0.2 ×10^7^ PFU/mL at 40 hpi), indicating that EtOH had no effect on virus progeny production. More than 100-fold reduction in virus particle production was observed at 16, 24 and 40 hpi following infection in presence of moxidectin or ivermectin **(Fig. 1C)**. At 60 hpi, ivermectin, showed antiviral activity for 2 experiments, yet there was a considerable variation between the other 2 experiments and therefore no significant antiviral effect was seen. In contrast, a moderate but consistent antiviral effect of moxidectin was seen at 60 hpi. At this time point the titer reduced from 9.9 □±□ 2.9 ×10^6^ PFU/mL (EtOH control) to 1.5 □±□ 1.0 ×10^6^ PFU/mL, which corresponds to a reduction of 84.8%. Taken together, a single 10 µM dose of both compounds controlled SARS-CoV-2 replication for at least 5 replication cycles. For moxidectin, a more consistent antiviral effect is observed lasting until 7 replication cycles.

### 3.2 Moxidectin and ivermectin interfere with SARS-CoV-2 replication

To delineate the mode-of-action of the drug, we first investigated whether moxidectin and ivermectin exhibit a direct virucidal effect. To this end, 10 µM of moxidectin, ivermectin or the corresponding amount of EtOH control was incubated with 2.5 ×10^5^ SARS-CoV-2 particles at 37 °C for 2 h and the infectious titer was determined by plaque assay. The highest concentration in the diluted sample corresponded to 1 µM compound and at this concentration no antiviral effect in Vero E6 cells was observed **(Fig. 1B)** and therefore the plaque assay can be used as readout. Importantly, no differences in viral titer were observed between the moxidectin or ivermectin-treated samples and the EtOH control **(Fig. 2A)**, indicating that these compounds do not exhibit virucidal activity to SARS-CoV-2 particles. Next, we performed a time-of-drug-addition assay. Here, 10 µM of moxidectin or ivermectin was administered either prior to virus inoculation, during virus inoculation, or post-virus inoculation **(Fig. 2B)**. Cells were infected with SARS-CoV-2 at an MOI of 1 and supernatants were collected at 8 hpi to determine progeny infectious virus particle production. Each treatment included a corresponding EtOH control. No effect on the viral titers were observed when the compounds were added prior to inoculation. For the ‘during’ condition, no significant effect was seen for moxidectin and a significant albeit limited effect was observed for ivermectin. A strong reduction in viral titer was seen when the compounds were added at 2 hpi **(Fig. 2C)**. For moxidectin, the infectious virus titer was reduced to 1.5 ± 1.0 ×10^3^ PFU/mL corresponding to a reduction of 98.6% and for ivermectin, the titer was reduced to 1.5 □ ± 0.8 ×10^2^ (99.9% reduction) when compared to the corresponding EtOH control. No significant effect was seen when moxidectin or ivermectin were added at 4 or 6 hpi when compared to EtOH control **(Fig. S2)**. Collectively, these results indicate that both moxidectin and ivermectin directly interfere with the viral infectious replication cycle in cells. This shows that the compounds either act at the early stages of RNA replication/translation (within 2-4 h post-infection) or that the compounds interfere with late stages of virus assembly/secretion (i.e. needs to be present for more than 4 h to exert its antiviral effect).

**Fig. 2.**
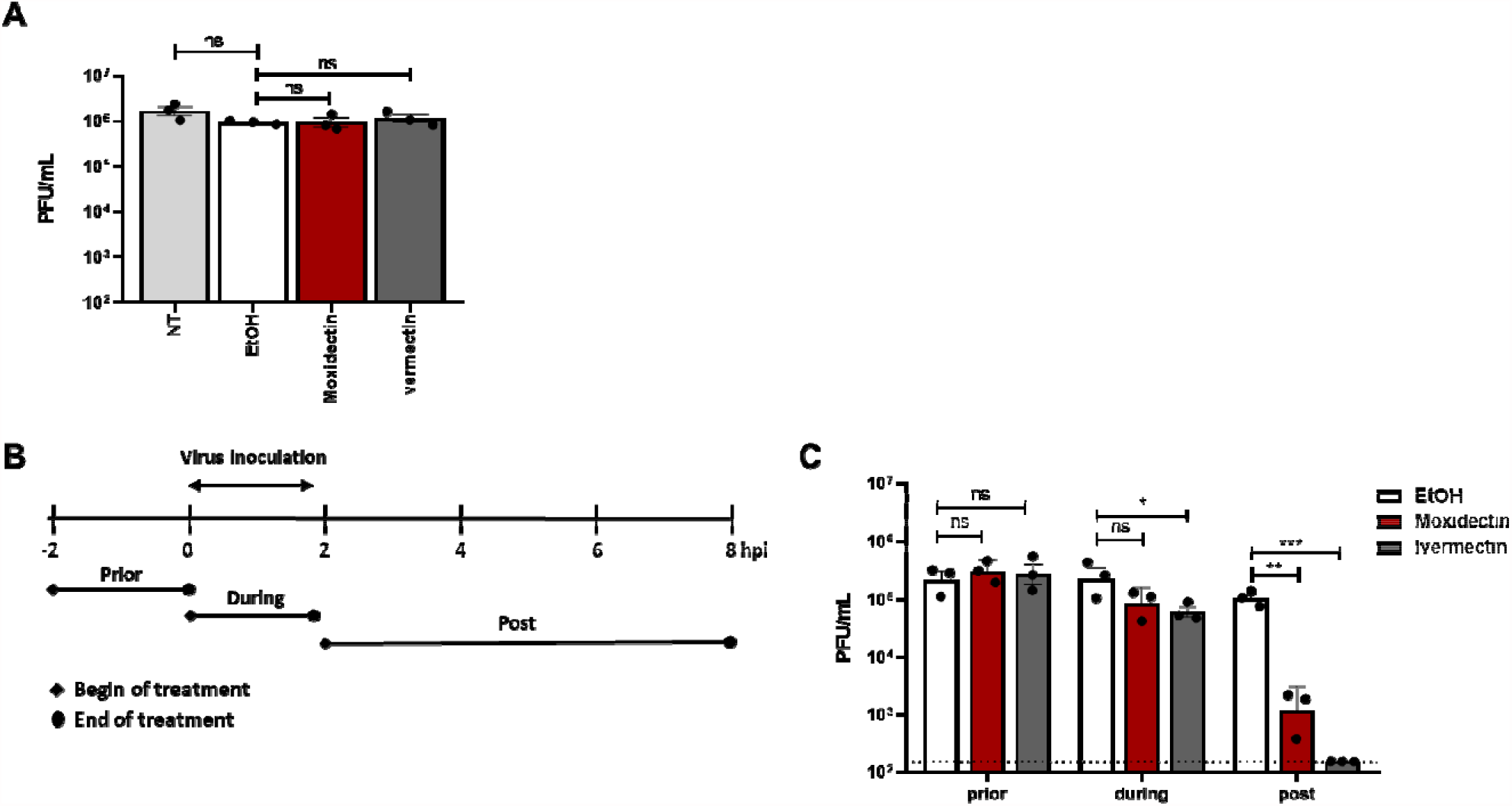
Moxidectin and ivermectin predominantly inhibit SARS-CoV-2 at post-infection conditions. **(A)** Virucidal effect of moxidectin and ivermectin on SARS-CoV-2. (**B**) Schematic summary of the time-of-drug-addition experiment. (**C**) Vero E6 cells were infected with SARS-CoV-2 at MOI 1 for 2 h following which the inoculum was removed. Cells were treated with moxidectin or ivermectin at a concentration of 10 µM or the corresponding volume of EtOH as depicted in (**B**). At 8 hpi, cell supernatants were harvested and the virus titer was determined via plaque assay. The dotted lines indicate the detection limit of plaque assay. Each dot represents data from a single independent experiment. Data are presented as mean ± SEM for three independent experiments. Statistical analysis was carried out by comparing treated samples with the EtOH control using Student’s t-test. *** p < 0.001, ** p< 0.01, and * p <0.05 and ns as non-significant

### 3.3 Moxidectin and ivermectin do not exhibit anti-SARS-CoV-2 activity in Calu-3 and primary human bronchial epithelial cells

We next sought to validate the antiviral potential of moxidectin and ivermectin in the human lung epithelial Calu-3 cells, cells which have previously been shown to support SARS-CoV-2 replication (Chu et al., 2020). Prior to infectivity assays, the cellular cytotoxicity of both compounds at 5 and 10 µM was determined using the MTS assay. At these concentrations, no cytotoxicity was observed **(Fig. S3)**. Accordingly, we proceeded by infecting Calu-3 cells with SARS-CoV-2 at MOI 1 in the presence or absence of 5 µM and 10 µM of moxidectin or ivermectin. In the absence of the drugs (NT condition), infection led to a release of on average, 1.5□±□2.3 ×10^4^ PFU/mL at 8 hpi **(Fig. 3A)**. A comparable titer was observed for the EtOH control (1.7□±□2.5 ×10^4^ PFU/mL). To our surprise, however, no significant reduction in viral titers was observed following infection in presence of moxidectin and ivermectin, indicating that at these experimental conditions, the compounds did not exhibit an antiviral effect in Calu-3 cells.

**Fig. 3.**
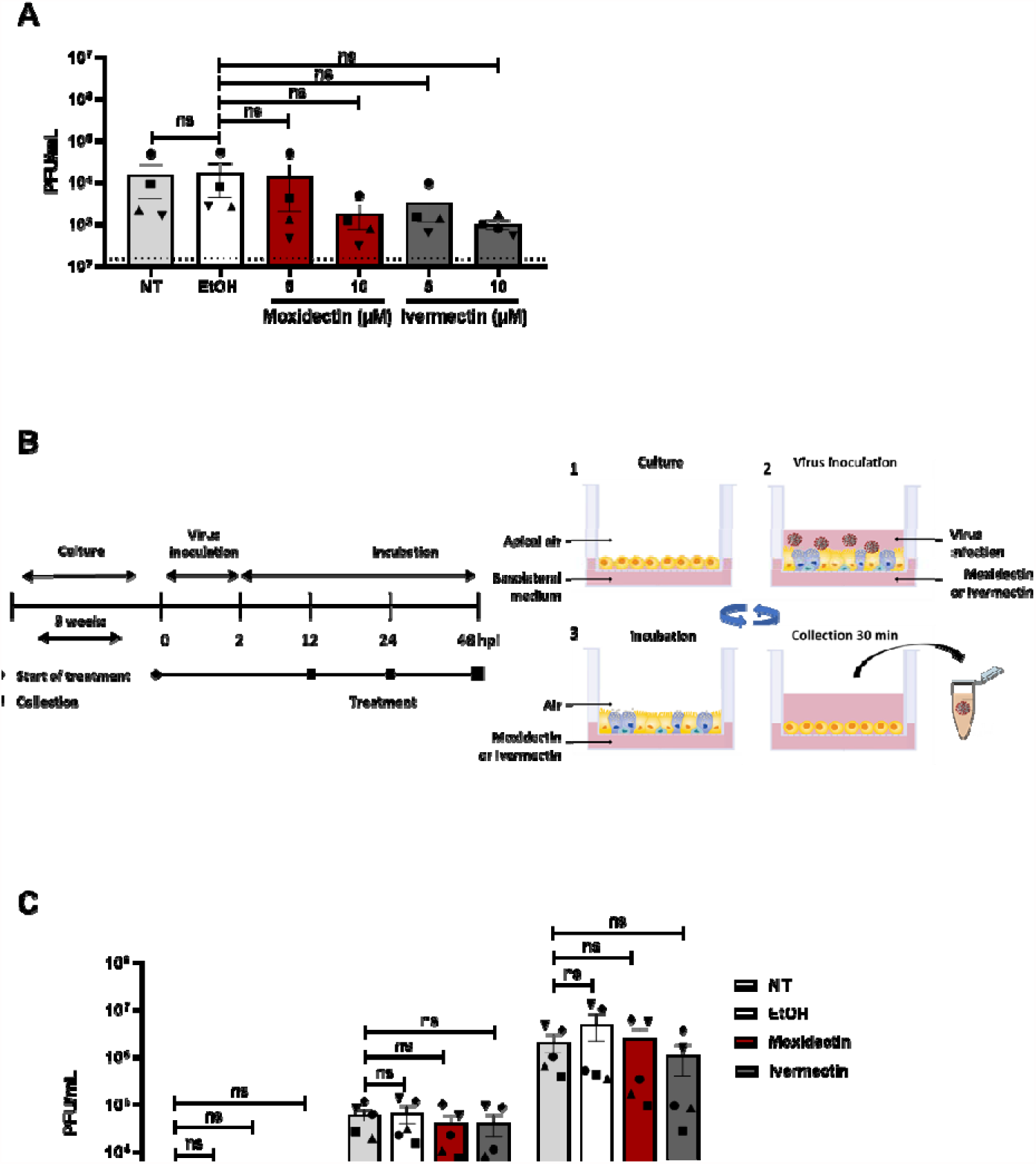
Moxidectin and ivermectin have no significant effect on SARS-CoV-2 infection in human-derived cell models. **(A)** Calu-3 cells were treated with 5 and 10 µM of moxidectin or ivermectin or the highest respective concentration of EtOH in presence of SARS-CoV-2 at MOI 1. At 8 hpi, cell supernatants were harvested and the number of produced infectious virus particles were determined by plaque assay. Each dot represents data from a single independent experiment. **(B)** Schematic representation of the experimental design. (1) Scheme of experiments in primary (PBECs). The cells were cultured on permeable inserts under Air Liquid Interface (ALI) conditions for ∼ 3 weeks. (2) Cells were inoculated with SARS-CoV-2 MOI 5 at the apical side of the insert and treated with 10 µM of moxidectin, ivermectin or the corresponding volume of EtOH at basolateral side until the collection time as shown in the schematics (3) Following virus inoculation, virus was removed by washing with media and the cells were left under ALI conditions until virus collection. (4) For final collection, cells were incubated with medium for 30 min and produced progeny virus was collected. After collection, cells were exposed to air again (3) until the next collection time point where steps 4 and 3 were repeated until the end of treatment. **(C)** The antiviral activity of moxidectin and ivermectin in PBECs. Dotted lines indicate the threshold of detection. Scheme adapted from STEMCELL Technologies (STEMCELL, 2021). Each dot represents data from a single independent experiment. Data is represented as mean ± SEM from five different donors. Student t-test was used to evaluate statistical differences and a p value ≤ 0.05 was considered significant with *p ≤ 0.05, **p ≤ 0.01 and ***p ≤ 0.001 and ns as non-significant.

As the human respiratory tract represents the primary site of virus infection (Jonsdottir and Dijkman, 2016), we decided to further verify the antiviral activity of moxidectin and ivermectin in a human-based cell system. Hereto, we used PBECs cultured under ALI culture system. Cells grown under ALI conditions undergo cellular differentiation, thus mimicking crucial physiological properties similar to that found *in vivo* (Cao et al., 2020; Jia et al., 2005; Sims et al., 2008). Prior to performing the antiviral assays in this model system, we first assessed the cytotoxicity of 10 µM moxidectin and ivermectin at 48 h of treatment using live-death staining and a LDH release assay. Both assays showed a viability above 90% compared to EtOH control **(Fig. S4)**. ALI-cultured PBECs were incubated with 10 µM of moxidectin or ivermectin at the basolateral side and infected with SARS-CoV-2 at MOI 5 at the apical side as indicated in the schematics **(Fig. 3B)**. Supernatants were harvested from the apical side at 12, 24 and 48 hpi and titers were determined using plaque assay. Importantly, despite the strong antiviral effect in Vero E6 cells, no significant antiviral activity was seen in the apical washes of moxidectin or ivermectin treated PBECs when compared to the EtOH control at all-time points tested **(Fig. 3C)**. Next to moxidectin and ivermectin, we also tested antiviral activity of resveratrol in the same donors and observed a potent antiviral effect in this model system thereby confirming the validity of the experimental settings (Ellen et al., 2020).

## 4. Discussion

We confirm previously published antiviral activity of ivermectin (Caly et al., 2020) in Vero E6 cells and showed an antiviral effect of moxidectin against SARS-CoV-2 in these cells. Further in-depth mode-of-action experiments revealed that moxidectin and ivermectin actively interfere with virus replication. Unfortunately, despite the promising antiviral properties of moxidectin and ivermectin in Vero E6 cells, the drugs did not show any antiviral activity in human airway-derived cell models. Collectively, the data show that the potency of moxidectin and ivermectin are species or cell-type specific and provide an explanation for the discordant findings in literature where positive effects were found in Vero cells, but no clinically relevant patient outcomes could be demonstrated in human studies.

To demonstrate which steps of the SARS-CoV-2 replication cycle are affected by moxidectin or ivermectin in Vero E6 cells, we performed a time-of-drug-addition experiments. The strongest effect was seen when moxidectin or ivermectin are added after the onset of infection suggesting the compounds do not interfere with processes related to viral entry like virus-cell binding, internalization or membrane fusion. We hypothesize that the compounds either directly interfere with early stages of RNA translation/replication or indirectly at late stages of the virus replication cycle (due to prolonged incubation). Other studies revealed that ivermectin blocks replication of HIV-1 and DENV viruses (Tay et al., 2013; Wagstaff et al., 2012), by binding and inhibiting cellular importin (IMP) α/β-mediated transport of viral proteins to the nucleus (Wagstaff et al., 2011; Wagstaff et al., 2012). Ivermectin has also been shown to inhibit NS3 helicase activity thereby inhibiting flavivirus replication (Mastrangelo et al., 2012). Our findings hint towards a mechanism where both drugs restrict SARS-CoV-2 replication by modulating cellular factors or processes, which control replication in Vero E6 cells but are absent in human-derived Calu-3 and PBECs.

Currently (April 29^th^, 2021), 65 clinical trials are registered that evaluate the efficacy of ivermectin as a prophylactic or therapeutic drug and 21 of these are completed (clinicaltrials.gov, 2021). Many of these trials have serious limitations such as small sample size, lack of binding, lack of pre-registration for some trials and therefore carries a risk of bias (Ahmed et al., 2021; Chaccour et al., 2021; Chowdhury et al., 2020; Elgazzar et al., 2020; Hashim et al., 2020; Okumuş et al., 2021; Podder et al., 2020; Rajter et al., 2021; Soto-Becerra et al., 2020; WHO, 2021b). Apart from the problem of bias, only five trials (April 29, 2021), directly compared ivermectin with standard of care and reported clinically crucial outcomes such as mortality (Gonzalez et al., 2021; Lopez-Medina et al., 2021; Mohan et al., 2021; Niaee et al., 2020; Ravikirti. et al., 2021). The WHO recently stated: “the current evidence of effect of ivermectin on mortality, mechanical ventilation, hospital admission and duration of hospitalization remains uncertain” (WHO, 2021b). Yet, many clinical trials are still ongoing. The observed lack of antiviral activity of ivermectin in the biologically relevant PBECs model reported here together with the poor pharmacological properties of ivermectin (Momekov and Momekova, 2020; Pena-Silva et al., 2020; Schmith et al., 2020) do not forecast a success of the ongoing clinical trials for its usage in COVID-19 patients.

SARS-CoV-2 can also infect several economically important livestock such as cats, dogs, minks, lions and tigers (Lam et al., 2020; McAloose et al., 2020; Oreshkova et al., 2020; Patterson et al., 2020; Sharun et al., 2020; Shi et al., 2020). Inhibition of virus replication in these animals could thus offer an effective strategy to control virus dissemination. Both drugs inhibited SARS-CoV-2 infection in African green monkey-derived kidney epithelial (Vero E6) cells. Since both the compounds have already been FDA-approved for the use in animals (Gonzalez Canga et al., 2009; Nolan and Lok, 2012), our results in Vero E6, warrant follow-up studies in relevant animal cell systems to assess their potential as antivirals in economically important livestock.

In conclusion, we elucidated that while both moxidectin and ivermectin exhibit antiviral activity in Vero E6, they do not reduce SARS-CoV-2 replication in human airway-derived cell models. While immortalized cells lines such as Vero E6 may be useful in a primary screening of inhibitors, it is important to determine true relevance of the inhibitors in a biological relevant model. In humans, airway epithelium is the main target for SARS-CoV-2 (de Melo et al., 2020; Leist et al., 2020) and therefore PBECs grown under ALI conditions, but not Vero cells, mimic crucial physiological properties similar to that found *in vivo* (Cao et al., 2020; Jia et al., 2005; Sims et al., 2008). We therefore advocate more rigorous antiviral drug testing in relevant systems prior to evaluating their efficacy in clinical trials.

## Acknowledgements

The authors thank dr. Y. Bhide for help in the design of experiments. Grant support was provided by University Medical Center of Groningen and Marie Skłodowska-Curie Cofund (713660). N.D.K, B.M.E, E.M.B, M.C.N, Y.S, I.A.R.Z. and J.M.S designed the experiments. N.D.K, B.M.E, E.M.B, B.T, D.P.I.P, H.H.E, D.G, L.A, executed the experiments. O.A.C and M.B selected recruited donors and sampled donors. N.D.K, E.M.B. I.A.R.Z and J.M.S wrote the manuscript. All authors edited the manuscript.

## Supporting information

**Fig. S1.**
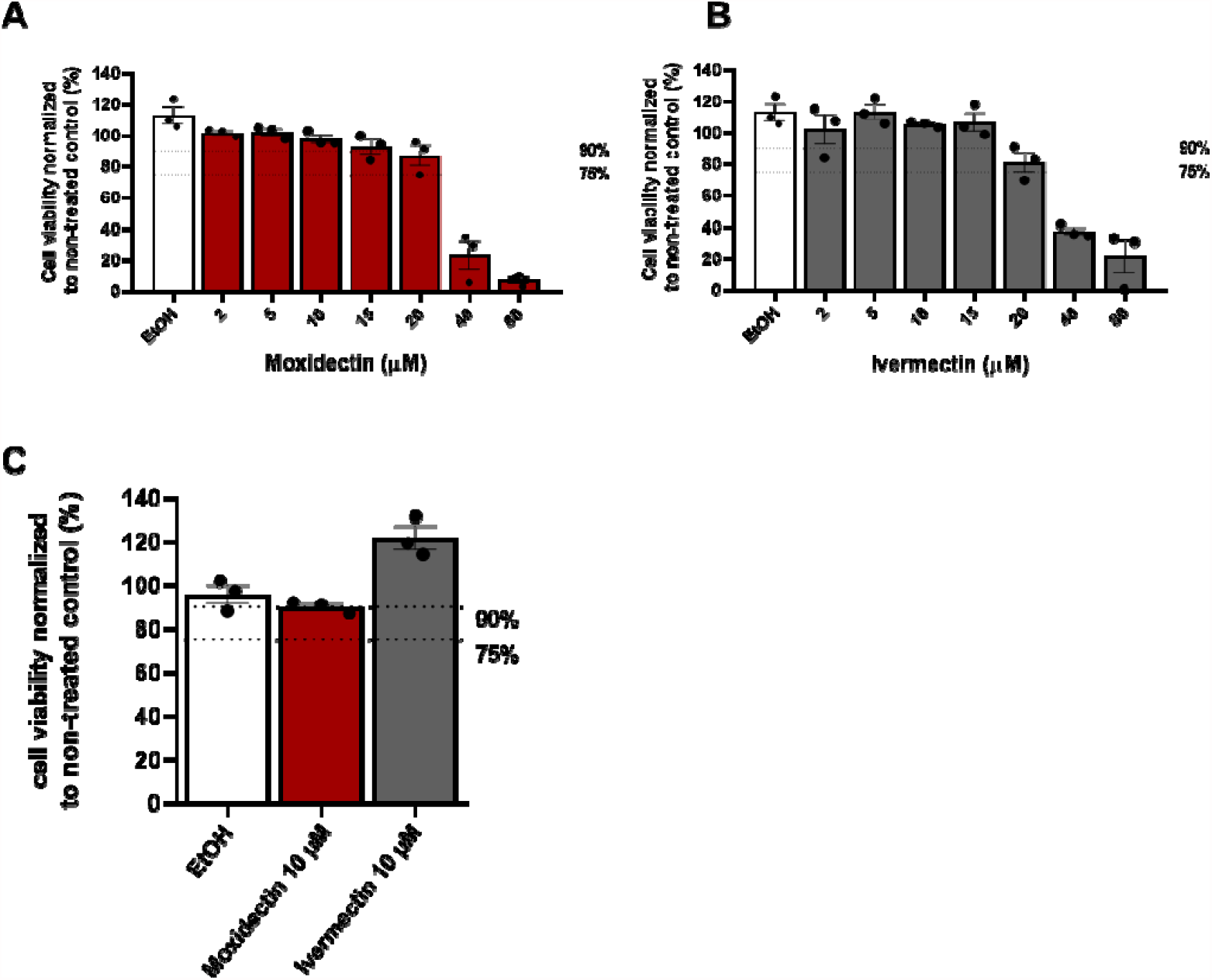
Cellular cytotoxicity of moxidectin and ivermectin in Vero E6 cells. Dose-dependent cytotoxicity of (**A**) moxidectin, (**B**) ivermectin in Vero E6 cells. Cells were incubated for 8 h with increasing concentrations of the compound or equivalent volume of EtOH corresponding to the highest concentration of compound **(C)** Vero E6 cells were incubated for prolonged incubation of 60 h with moxidectin (10 µM), ivermectin (10 µM) or equivalent volumes of EtOH. Cellular cytotoxicity was determined by MTS assay. Cell survival is expressed as percentage compared to non-treated control. Each experiment was carried out in triplicate. Each dot represents data from a single independent experiment. Data are represented as mean ± SEM of at least three independent experiments.

**Fig. S2.**
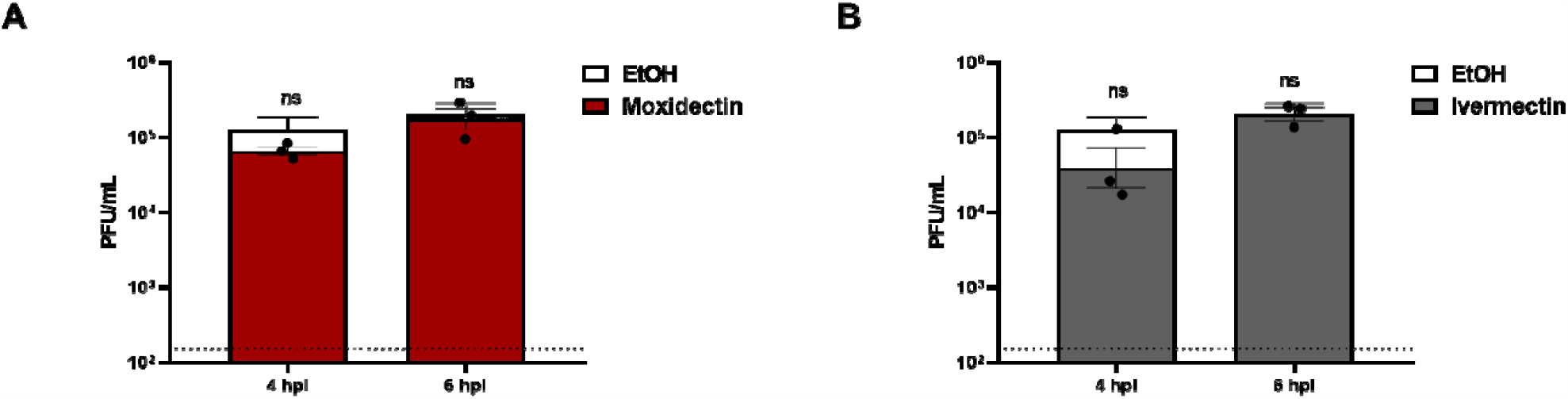
Effect of moxidectin and ivermectin upon addition late in replication cycle of SARS-CoV-2. Vero E6 cells were infected with SARS-CoV-2 at MOI 1 for 2 h following which the inoculum was removed. Cells were treated with **(A)** moxidectin or **(B)** ivermectin at a concentration of 10 µM or the corresponding volume of EtOH at 4 or 6 hpi. At 8 hpi, cell supernatants were harvested and the virus titer was determined via plaque assay. The dotted lines indicate the detection limit of plaque assay. Each dot represents data from a single independent experiment. Data are represented as mean ± SEM of at least three independent experiments. Statistical analysis was carried out by comparing treated samples with the EtOH control using Student’s t-test *** p < 0.001, ** p< 0.01, * p <0.05 and ns as non-significant.

**Fig. S3.**
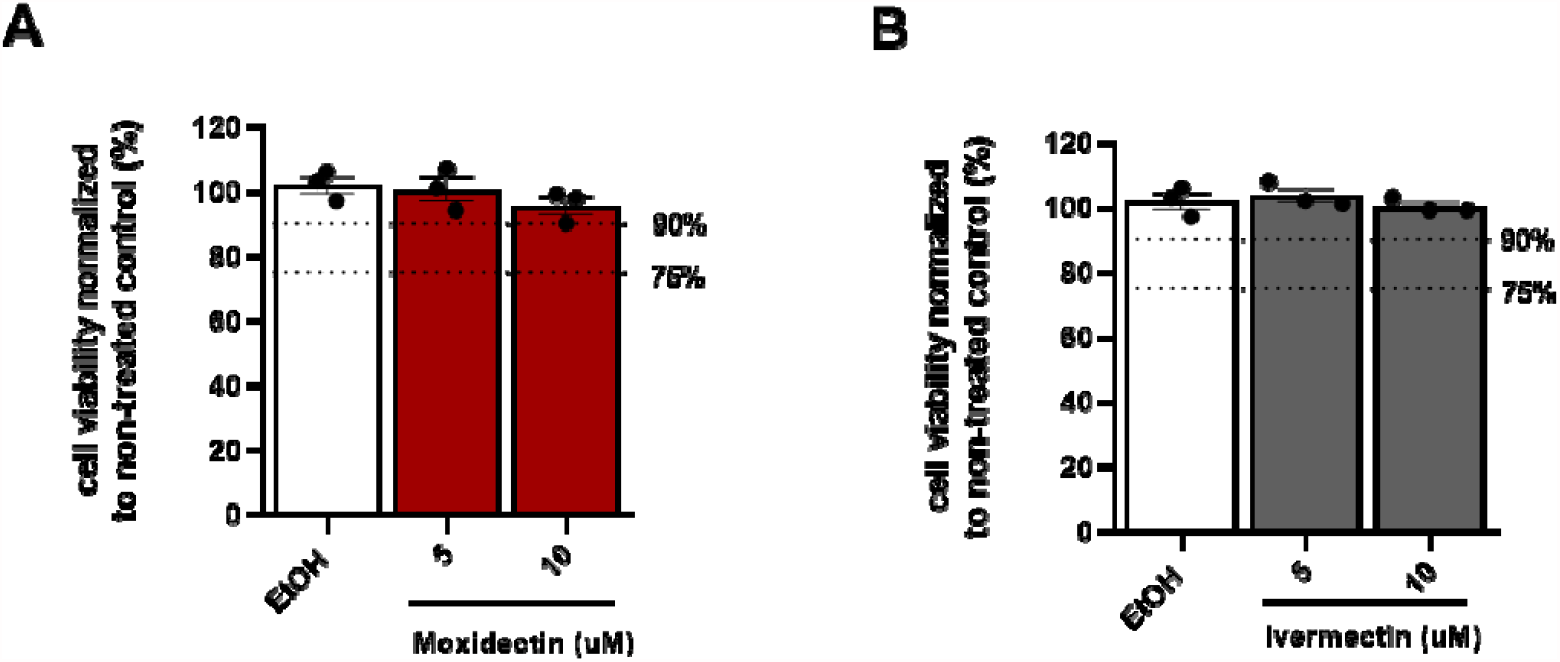
Cellular cytotoxicity of moxidectin and ivermectin in Calu-3 cells. Calu-3 cells were incubated with moxidectin (5 and 10 µM), ivermectin (5 and 10 µM) or equivalent volumes of EtOH corresponding to the highest used concentration, respectively. Cellular cytotoxicity was determined by MTS assay. Cell survival is expressed as percentage compared to non-treated control. Each dot represents data from a single independent experiment. Data are represented as mean ± SEM of at least three independent experiments.

**Fig. S4.**
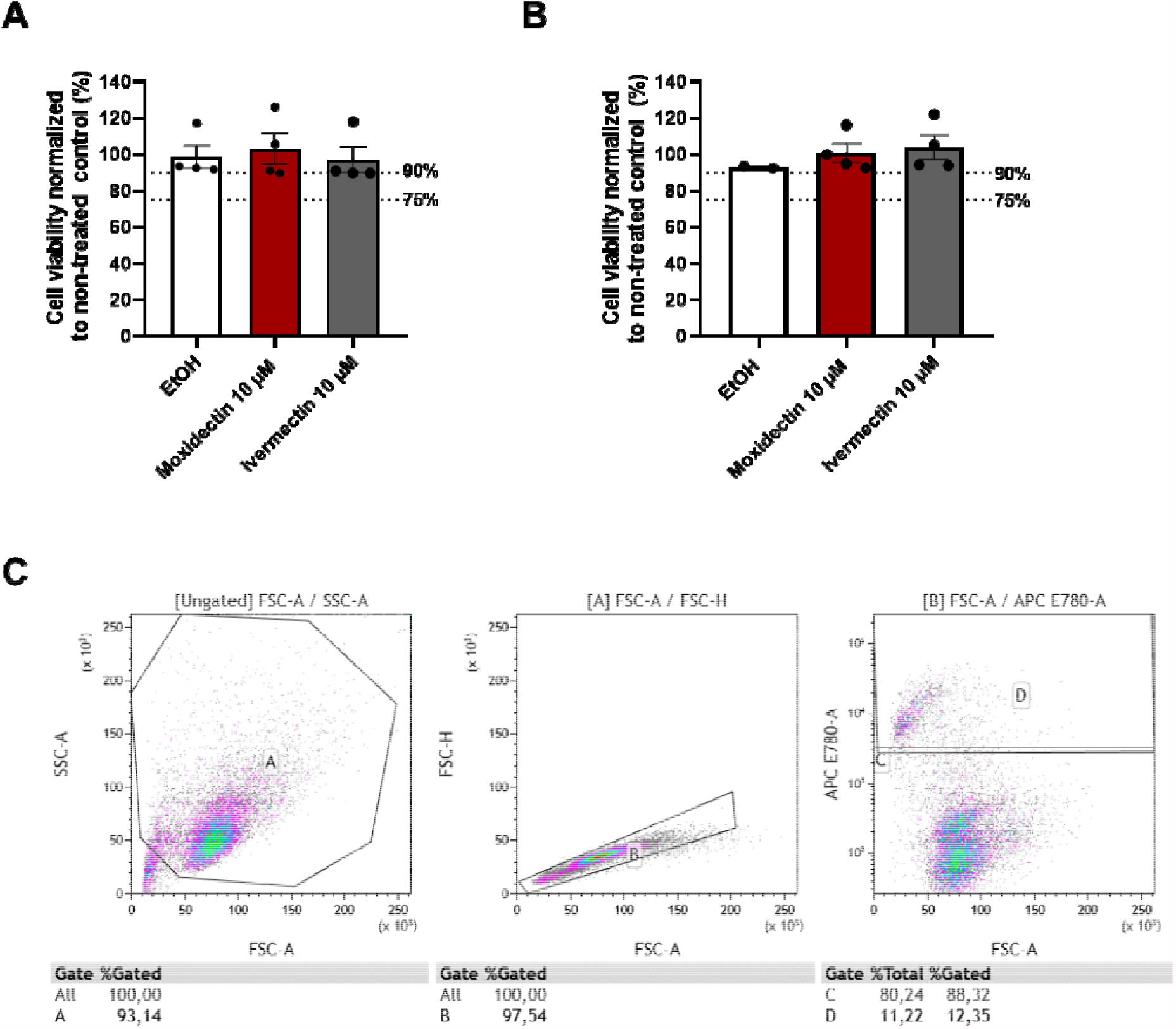
Cellular cytotoxicity of moxidectin and ivermectin in PBECs. PBECs were either left untreated (NT) or incubated with 10 µM moxidectin, 10 µM ivermectin or the corresponding volume of EtOH at basolateral side for 8 hr. Cell viability was determined by flow cytometry using **(A)** viability dye and by **(B)** LDH assay. **(C)** Gating strategy used for flow cytometry. Each dot represents data from a single independent experiment. Data are represented as mean ± SEM of at least three independent experiments.

## References

Ahmed, S., Karim, M.M., Ross, A.G., Hossain, M.S., Clemens, J.D., Sumiya, M.K., Phru, C.S., Rahman, M., Zaman, K., Somani, J., Yasmin, R., Hasnat, M.A., Kabir, A., Aziz, A.B., Khan, W.A., 2021. A five-day course of ivermectin for the treatment of COVID-19 may reduce the duration of illness. Int J Infect Dis 103, 214–216.

Bray, M., Rayner, C., Noel, F., Jans, D., Wagstaff, K., 2020. Ivermectin and COVID-19: A report in Antiviral Research, widespread interest, an FDA warning, two letters to the editor and the authors’ responses. Antiviral Res 178, 104805.

Caly, L., Druce, J.D., Catton, M.G., Jans, D.A., Wagstaff, K.M., 2020. The FDA-approved drug ivermectin inhibits the replication of SARS-CoV-2 in vitro. Antiviral Res 178, 104787.

Cao, X., Coyle, J.P., Xiong, R., Wang, Y., Heflich, R.H., Ren, B., Gwinn, W.M., Hayden, P., Rojanasakul, L., 2020. Invited review: human air-liquid-interface organotypic airway tissue models derived from primary tracheobronchial epithelial cells-overview and perspectives. In Vitro Cell Dev Biol Anim.

Chaccour, C., Casellas, A., Blanco-Di Matteo, A., Pineda, I., Fernandez-Montero, A., Ruiz-Castillo, P., Richardson, M.A., Rodriguez-Mateos, M., Jordan-Iborra, C., Brew, J., Carmona-Torre, F., Giraldez, M., Laso, E., Gabaldon-Figueira, J.C., Dobano, C., Moncunill, G., Yuste, J.R., Del Pozo, J.L., Rabinovich, N.R., Schoning, V., Hammann, F., Reina, G., Sadaba, B., Fernandez-Alonso, M., 2021. The effect of early treatment with ivermectin on viral load, symptoms and humoral response in patients with non-severe COVID-19: A pilot, double-blind, placebo-controlled, randomized clinical trial. EClinicalMedicine, 100720.

Chan, W., He, B., Wang, X., He, M.-L., 2020. Pandemic COVID-19: current status and challenges of antiviral therapies. Genes & Diseases.

Chowdhury, A., Shahbaz, M., Karim, M.R., 2020. A Randomized Trial of Ivermectin-Doxycycline and Hydroxychloroquine-Azithromycin therapy on COVID19 patients. Preprint Research Square.

Chu, H., Chan, J.F., Yuen, T.T., Shuai, H., Yuan, S., Wang, Y., Hu, B., Yip, C.C., Tsang, J.O., Huang, X., Chai, Y., Yang, D., Hou, Y., Chik, K.K., Zhang, X., Fung, A.Y., Tsoi, H.W., Cai, J.P., Chan, W.M., Ip, J.D., Chu, A.W., Zhou, J., Lung, D.C., Kok, K.H., To, K.K., Tsang, O.T., Chan, K.H., Yuen, K.Y., 2020. Comparative tropism, replication kinetics, and cell damage profiling of SARS-CoV-2 and SARS-CoV with implications for clinical manifestations, transmissibility, and laboratory studies of COVID-19: an observational study. Lancet Microbe 1, e14-e23.

clinicaltrials.gov, 2021. Ivermectin and COVID-19.

Cobb, R., Boeckh, A., 2009. Moxidectin: a review of chemistry, pharmacokinetics and use in horses. Parasit Vectors 2 Suppl 2, S5.

Consortium, W.S.T., Pan, H., Peto, R., Henao-Restrepo, A.M., Preziosi, M.P., Sathiyamoorthy, V., Abdool Karim, Q., Alejandria, M.M., Hernandez Garcia, C., Kieny, M.P., Malekzadeh, R., Murthy, S., Reddy, K.S., Roses Periago, M., Abi Hanna, P., Ader, F., Al-Bader, A.M., Alhasawi, A., Allum, E., Alotaibi, A., Alvarez-Moreno, C.A., Appadoo, S., Asiri, A., Aukrust, P., Barratt-Due, A., Bellani, S., Branca, M., Cappel-Porter, H.B.C., Cerrato, N., Chow, T.S., Como, N., Eustace, J., Garcia, P.J., Godbole, S., Gotuzzo, E., Griskevicius, L., Hamra, R., Hassan, M., Hassany, M., Hutton, D., Irmansyah, I., Jancoriene, L., Kirwan, J., Kumar, S., Lennon, P., Lopardo, G., Lydon, P., Magrini, N., Maguire, T., Manevska, S., Manuel, O., McGinty, S., Medina, M.T., Mesa Rubio, M.L., Miranda-Montoya, M.C., Nel, J., Nunes, E.P., Perola, M., Portoles, A., Rasmin, M.R., Raza, A., Rees, H., Reges, P.P.S., Rogers, C.A., Salami, K., Salvadori, M.I., Sinani, N., Sterne, J.A.C., Stevanovikj, M., Tacconelli, E., Tikkinen, K.A.O., Trelle, S., Zaid, H., Rottingen, J.A., Swaminathan, S., 2021. Repurposed Antiviral Drugs for Covid-19 -Interim WHO Solidarity Trial Results. N Engl J Med 384, 497–511.

de Melo, B.A.G., Benincasa, J.C., Cruz, E.M., Maricato, J.T., Porcionatto, M.A., 2020. 3D culture models to study SARS-CoV-2 infectivity and antiviral candidates: From spheroids to bioprinting. Biomed J.

Elgazzar, A., Hany, B., Youssef, S.A., Hany, B., Hafez, M., Moussa, H., 2020. Efficacy and Safety of Ivermectin for Treatment and prophylaxis of COVID-19 Pandemic. Research square Preprint.

Ellen, B.M.T., Kumar, N.D., Bouma, E.M., Troost, B., Van de Pol, D.P.I., van der Ende-Metselaar, H.H., Apperloo, L., Gosliga, D.V., van den Berge, M., Nawijn, M.C., van der Voort, P.H.J., Moser, J., Rodenhuis-Zybert, I.A., Smit, J.M., 2020. Resveratrol And Pterostilbene Potently Inhibit SARS-CoV-2 Replication In Vitro. bioRxiv preprint.

EMA, 2009. Ivermectin.

EMA, 2017. Moxidectin-containing veterinary medicines used in cattle, sheep and horses

Gonzalez Canga, A., Sahagun Prieto, A.M., Jose Diez Liebana, M., Martinez, N.F., Vega, M.S., Vieitez, J.J., 2009. The pharmacokinetics and metabolism of ivermectin in domestic animal species. Vet J 179, 25–37.

Gonzalez, J.L.B., Gámez, M.G., Enciso, E.A.M., Maldonado, R.J.E., Palacios, D.H., Campos, S.D., Robles, I.O., Guzmán, M.J.M., Díaz, A.L.G., Peña, C.M.G., Medina, L.M., Colin, V.A.M., Manuel, A.G.J., 2021. Efficacy and safety of Ivermectin and Hydroxychloroquine in patients with severe COVID-19. A randomized controlled trial. medRxiv Preprint.

Gotz, V., Magar, L., Dornfeld, D., Giese, S., Pohlmann, A., Hoper, D., Kong, B.W., Jans, D.A., Beer, M., Haller, O., Schwemmle, M., 2016. Influenza A viruses escape from MxA restriction at the expense of efficient nuclear vRNP import. Sci Rep 6, 23138.

Hashim, H.A., Maulood, M., Rasheed, A.M., Fatak, D.F., Kabah, K.K., S, A.A., 2020. Controlled randomized clinical trial on using Ivermectin with Doxycycline for treating COVID-19 patients in Baghdad, Iraq. medRxiv Preprint.

Heijink, I.H., Kies, P.M., Kauffman, H.F., Postma, D.S., van Oosterhout, A.J., Vellenga, E., 2007. Down-regulation of E-cadherin in human bronchial epithelial cells leads to epidermal growth factor receptor-dependent Th2 cell-promoting activity. J Immunol 178, 7678–7685.

Heijink, I.H., Postma, D.S., Noordhoek, J.A., Broekema, M., Kapus, A., 2010. House dust mite-promoted epithelial-to-mesenchymal transition in human bronchial epithelium. Am J Respir Cell Mol Biol 42, 69–79.

Jia, H.P., Look, D.C., Shi, L., Hickey, M., Pewe, L., Netland, J., Farzan, M., Wohlford-Lenane, C., Perlman, S., McCray, P.B., Jr., 2005. ACE2 receptor expression and severe acute respiratory syndrome coronavirus infection depend on differentiation of human airway epithelia. J Virol 79, 14614–14621.

Jonsdottir, H.R., Dijkman, R., 2016. Coronaviruses and the human airway: a universal system for virus-host interaction studies. Virol J 13, 24.

Ketkar, H., Yang, L., Wormser, G.P., Wang, P., 2019. Lack of efficacy of ivermectin for prevention of a lethal Zika virus infection in a murine system. Diagn Microbiol Infect Dis 95, 38–40.

Lam, S.D., Bordin, N., Waman, V.P., Scholes, H.M., Ashford, P., Sen, N., van Dorp, L., Rauer, C., Dawson, N.L., Pang, C.S.M., Abbasian, M., Sillitoe, I., Edwards, S.J.L., Fraternali, F., Lees, J.G., Santini, J.M., Orengo, C.A., 2020. SARS-CoV-2 spike protein predicted to form complexes with host receptor protein orthologues from a broad range of mammals. Sci Rep 10, 16471.

Leist, S.R., Schafer, A., Martinez, D.R., 2020. Cell and animal models of SARS-CoV-2 pathogenesis and immunity. Dis Model Mech 13.

Lopez-Medina, E., Lopez, P., Hurtado, I.C., Davalos, D.M., Ramirez, O., Martinez, E., Diazgranados, J.A., Onate, J.M., Chavarriaga, H., Herrera, S., Parra, B., Libreros, G., Jaramillo, R., Avendano, A.C., Toro, D.F., Torres, M., Lesmes, M.C., Rios, C.A., Caicedo, I., 2021. Effect of Ivermectin on Time to Resolution of Symptoms Among Adults With Mild COVID-19: A Randomized Clinical Trial. JAMA 325, 1426–1435.

Lundberg, L., Pinkham, C., Baer, A., Amaya, M., Narayanan, A., Wagstaff, K.M., Jans, D.A., Kehn-Hall, K., 2013. Nuclear import and export inhibitors alter capsid protein distribution in mammalian cells and reduce Venezuelan Equine Encephalitis Virus replication. Antiviral Res 100, 662–672.

Makhani, L., Khatib, A., Corbeil, A., Kariyawasam, R., Raheel, H., Clarke, S., Challa, P., Hagopian, E., Chakrabarti, S., Schwartz, K.L., Boggild, A.K., 2019. 2018 in review: five hot topics in tropical medicine. Trop Dis Travel Med Vaccines 5, 5.

Mastrangelo, E., Pezzullo, M., De Burghgraeve, T., Kaptein, S., Pastorino, B., Dallmeier, K., de Lamballerie, X., Neyts, J., Hanson, A.M., Frick, D.N., Bolognesi, M., Milani, M., 2012. Ivermectin is a potent inhibitor of flavivirus replication specifically targeting NS3 helicase activity: new prospects for an old drug. J Antimicrob Chemother 67, 1884–1894.

Matsuyama, S., Nao, N., Shirato, K., Kawase, M., Saito, S., Takayama, I., Nagata, N., Sekizuka, T., Katoh, H., Kato, F., Sakata, M., Tahara, M., Kutsuna, S., Ohmagari, N., Kuroda, M., Suzuki, T., Kageyama, T., Takeda, M., 2020. Enhanced isolation of SARS-CoV-2 by TMPRSS2-expressing cells. Proc Natl Acad Sci U S A 117, 7001–7003.

McAloose, D., Laverack, M., Wang, L., Killian, M.L., Caserta, L.C., Yuan, F., Mitchell, P.K., Queen, K., Mauldin, M.R., Cronk, B.D., Bartlett, S.L., Sykes, J.M., Zec, S., Stokol, T., Ingerman, K., Delaney, M.A., Fredrickson, R., Ivancic, M., Jenkins-Moore, M., Mozingo, K., Franzen, K., Bergeson, N.H., Goodman, L., Wang, H., Fang, Y., Olmstead, C., McCann, C., Thomas, P., Goodrich, E., Elvinger, F., Smith, D.C., Tong, S., Slavinski, S., Calle, P.P., Terio, K., Torchetti, M.K., Diel, D.G., 2020. From People to Panthera: Natural SARS-CoV-2 Infection in Tigers and Lions at the Bronx Zoo. mBio 11.

Milton, P., Hamley, J.I.D., Walker, M., Basanez, M.G., 2020. Moxidectin: an oral treatment for human onchocerciasis. Expert Rev Anti Infect Ther, 1–15.

Mohan, A., Tiwari, P., Suri, T., Mittal, S., Patel, A., Jain, A.T.V., Das, U.A., Bopanna, T.K., Pandey, R.M., Shelke, S., Singh, A.R., Bhatnagar, S., Masih, S., Mahajan, S., Dwivedi, T., Sahoo, B., Pandit, A., Bhopale, S., Vig, S., Gupta, R., Madan, K., Hadda, V., Gupta, N., Garg, R., Meena, V.P., Guleria, R., 2021. Ivermectin in mild and moderate COVID-19 (RIVET-COV): a randomized, placebo-controlled trial. Research square Preprint.

Momekov, G., Momekova, D., 2020. Ivermectin as a potential COVID-19 treatment from the pharmacokinetic point of view: antiviral levels are not likely attainable with known dosing regimens. Biotechnology & Biotechnological Equipment 34, 469–474.

Niaee, M.S., Gheibi, N., Namdar, P., Allami, A., Zolghadr, L., Javadi, A., Karampour, A., Varnaseri, M., Bizhani, B., Cheraghi, F., Naderi, Y., Amini, F., Karamyan, M., Yadyad, M.J., Jamshidian, R., 2020. Ivermectin as an adjunct treatment for hospitalized adult COVID-19 patients: A randomized multi-center clinical trial. Research square Preprint. NIH, 2021. Therapeutic Management of Adults With COVID-19.

Nolan, T.J., Lok, J.B., 2012. Macrocyclic lactones in the treatment and control of parasitism in small companion animals. Curr Pharm Biotechnol 13, 1078–1094.

Ogando, N.S., Dalebout, T.J., Zevenhoven-Dobbe, J.C., Limpens, R., van der Meer, Y., Caly, L., Druce, J., de Vries, J.J.C., Kikkert, M., Barcena, M., Sidorov, I., Snijder, E.J., 2020. SARS-coronavirus-2 replication in Vero E6 cells: replication kinetics, rapid adaptation and cytopathology. J Gen Virol 101, 925–940.

Okumuş, N., Demirtürk, N., Çetinkaya, R.A., Güner, R., Avci, I.Y., Orhan, S., Konya, P., şaylan, B., Karalezli, A., Yamanel, L., Kayaaslan, B., Yılmaz, G., Savaşçı, U., Eser, F., Taşkin, G., 2021. Evaluation of the Effectiveness and Safety of Adding Ivermectin to Treatment in Severe COVID-19 Patients. Research square Preprint.

Opoku, N.O., Bakajika, D.K., Kanza, E.M., Howard, H., Mambandu, G.L., Nyathirombo, A., Nigo, M.M., Kasonia, K., Masembe, S.L., Mumbere, M., Kataliko, K., Larbelee, J.P., Kpawor, M., Bolay, K.M., Bolay, F., Asare, S., Attah, S.K., Olipoh, G., Vaillant, M., Halleux, C.M., Kuesel, A.C., 2018. Single dose moxidectin versus ivermectin for Onchocerca volvulus infection in Ghana, Liberia, and the Democratic Republic of the Congo: a randomised, controlled, double-blind phase 3 trial. Lancet 392, 1207–1216.

Oreshkova, N., Molenaar, R.J., Vreman, S., Harders, F., Oude Munnink, B.B., Hakze-van der Honing, R.W., Gerhards, N., Tolsma, P., Bouwstra, R., Sikkema, R.S., Tacken, M.G., de Rooij, M.M., Weesendorp, E., Engelsma, M.Y., Bruschke, C.J., Smit, L.A., Koopmans, M., van der Poel, W.H., Stegeman, A., 2020. SARS-CoV-2 infection in farmed minks, the Netherlands, April and May 2020. Euro Surveill 25.

Patterson, E.I., Elia, G., Grassi, A., Giordano, A., Desario, C., Medardo, M., Smith, S.L., Anderson, E.R., Prince, T., Patterson, G.T., Lorusso, E., Lucente, M.S., Lanave, G., Lauzi, S., Bonfanti, U., Stranieri, A., Martella, V., Solari Basano, F., Barrs, V.R., Radford, A.D., Agrimi, U., Hughes, G.L., Paltrinieri, S., Decaro, N., 2020. Evidence of exposure to SARS-CoV-2 in cats and dogs from households in Italy. Nat Commun 11, 6231.

Pena-Silva, R., Duffull, S.B., Steer, A.C., Jaramillo-Rincon, S.X., Gwee, A., Zhu, X., 2020. Pharmacokinetic considerations on the repurposing of ivermectin for treatment of COVID-19. Br J Clin Pharmacol.

Podder, C.S., Chowdhury, N., Sina, M.I., Haque, W.M.M.U., 2020. Outcome of ivermectin treated mild to moderate COVID-19 cases: a single-centre, open-label, randomised controlled study. IMC Journal of Medical Science 14, 1–8.

Prichard, R., Menez, C., Lespine, A., 2012. Moxidectin and the avermectins: Consanguinity but not identity. Int J Parasitol Drugs Drug Resist 2, 134–153.

Prichard, R.K., Geary, T.G., 2019. Perspectives on the utility of moxidectin for the control of parasitic nematodes in the face of developing anthelmintic resistance. Int J Parasitol Drugs Drug Resist 10, 69–83.

Rajter, J.C., Sherman, M.S., Fatteh, N., Vogel, F., Sacks, J., Rajter, J.J., 2021. Use of Ivermectin Is Associated With Lower Mortality in Hospitalized Patients With Coronavirus Disease 2019: The Ivermectin in COVID Nineteen Study. Chest 159, 85–92.

Ravikirti Roy, R., Pattadar, C., Raj, R., Agarwal, N., Biswas, B., Majhi, P.M., Rai, D.K., Shyama. Kumar, A., Sarfaraz, A., 2021. Ivermectin as a potential treatment for mild to moderate COVID-19 – A double blind randomized placebo-controlled trial. MedRxiv Preprint.

Schmith, V.D., Zhou, J.J., Lohmer, L.R.L., 2020. The Approved Dose of Ivermectin Alone is not the Ideal Dose for the Treatment of COVID-19. Clin Pharmacol Ther 108, 762–765.

Sharun, K., Tiwari, R., Natesan, S., Dhama, K., 2020. SARS-CoV-2 infection in farmed minks, associated zoonotic concerns, and importance of the One Health approach during the ongoing COVID-19 pandemic. Vet Q, 1–14.

Shi, J., Wen, Z., Zhong, G., Yang, H., Wang, C., Huang, B., Liu, R., He, X., Shuai, L., Sun, Z., Zhao, Y., Liu, P., Liang, L., Cui, P., Wang, J., Zhang, X., Guan, Y., Tan, W., Wu, G., Chen, H., Bu, Z., 2020. Susceptibility of ferrets, cats, dogs, and other domesticated animals to SARS-coronavirus 2. Science 368, 1016–1020.

Siemieniuk, R., Rochwerg, B., Agoritsas, T., Lamontagne, F., Leo, Y.S., Macdonald, H., Agarwal, A., Zeng, L., Lytvyn, L., Appiah, J.A., Amin, W., Arabi, Y., Blumberg, L., Burhan, E., Bausch, F.J., Calfee, C.S., Cao, B., Cecconi, M., Chanda, D., Cooke, G., Du, B., Dunning, J., Geduld, H., Gee, P., Hashimi, M., Hui, D.S., Kabra, S., Kanda, S., Kawano-Dourado, L., Kim, Y.J., Kissoon, N., Kwizera, A., Laake, J.H., Machado, F.R., Mahaka, I., Manai, H., Mino, G., Nsutedu, E., Pshenichnaya, N., Qadir, N., Sabzwari, S., Sarin, R., Sharland, M., Shen, Y., Sri Ranganathan, S., Souza, J., Ugarte, S., Venkatapuram, S., Quoc Dat, V., Vuyiseka, D., Stegemann, M., Wijewickrama, A., Maguire, B., Zeraatkar, D., Bartoszko, J., Ge, L., Brignardello-Petersen, R., Owen, A., Guyatt, G., Diaz, J., Jacobs, M., Vandvik, P.O., 2020. A living WHO guideline on drugs for covid-19. BMJ 370, m3379.

Sims, A.C., Burkett, S.E., Yount, B., Pickles, R.J., 2008. SARS-CoV replication and pathogenesis in an in vitro model of the human conducting airway epithelium. Virus Res 133, 33–44.

Sohrabi, C., Alsafi, Z., O’Neill, N., Khan, M., Kerwan, A., Al-Jabir, A., Iosifidis, C., Agha, R., 2020. World Health Organization declares global emergency: A review of the 2019 novel coronavirus (COVID-19). Int J Surg 76, 71–76.

Soto-Becerra, P., Culquichicón, C., Hurtado-Roca, Y., Araujo-Castillo, R.V., 2020. Real-world effectiveness of hydroxychloroquine, azithromycin, and ivermectin among hospitalized COVID-19 patients: results of a target trial emulation using observational data from a nationwide healthcare system in Peru. medRxiv Preprint.

Stemcell, T., 2021. Air-Liquid Interface Culture for Respiratory Research

Tay, M.Y., Fraser, J.E., Chan, W.K., Moreland, N.J., Rathore, A.P., Wang, C., Vasudevan, S.G., Jans, D.A., 2013. Nuclear localization of dengue virus (DENV) 1-4 non-structural protein 5; protection against all 4 DENV serotypes by the inhibitor Ivermectin. Antiviral Res 99, 301–306.

Varghese, F.S., Kaukinen, P., Glasker, S., Bespalov, M., Hanski, L., Wennerberg, K., Kummerer, B.M., Ahola, T., 2016. Discovery of berberine, abamectin and ivermectin as antivirals against chikungunya and other alphaviruses. Antiviral Res 126, 117–124.

Vieira Braga, F.A., Kar, G., Berg, M., Carpaij, O.A., Polanski, K., Simon, L.M., Brouwer, S., Gomes, T., Hesse, L., Jiang, J., Fasouli, E.S., Efremova, M., Vento-Tormo, R., Talavera-Lopez, C., Jonker, M.R., Affleck, K., Palit, S., Strzelecka, P.M., Firth, H.V., Mahbubani, K.T., Cvejic, A., Meyer, K.B., Saeb-Parsy, K., Luinge, M., Brandsma, C.A., Timens, W., Angelidis, I., Strunz, M., Koppelman, G.H., van Oosterhout, A.J., Schiller, H.B., Theis, F.J., van den Berge, M., Nawijn, M.C., Teichmann, S.A., 2019. A cellular census of human lungs identifies novel cell states in health and in asthma. Nat Med 25, 1153–1163.

Wagstaff, K.M., Rawlinson, S.M., Hearps, A.C., Jans, D.A., 2011. An AlphaScreen(R)-based assay for high-throughput screening for specific inhibitors of nuclear import. J Biomol Screen 16, 192–200.

Wagstaff, K.M., Sivakumaran, H., Heaton, S.M., Harrich, D., Jans, D.A., 2012. Ivermectin is a specific inhibitor of importin alpha/beta-mediated nuclear import able to inhibit replication of HIV-1 and dengue virus. Biochem J 443, 851–856.

WHO, 2021a. Coronavirus disease (COVID-19) pandemic.

WHO, 2021b. Therapeutics and COVID-19 : living guidelines.

Yamasmith, E., Avirutnan, P., Mairiang, D., Tanrumluk, S., Suputtamongkol, Y., A-hamad Saleh-arong, F., Angkasekwinai, N., Wongsawa, E., Fongsri, U., 2018. Efficacy and safety of ivermectin against dengue infection: a phase III, randomized, double-blind, placebo-controlled trial, 34th Annual Meeting the Royal College of Physicians of Thailand. Internal Medicine and One Health, Chonburi, Thailand.

Zhang, J., Xie, B., Hashimoto, K., 2020. Current status of potential therapeutic candidates for the COVID-19 crisis. Brain Behav Immun 87, 59–73.

Zhou, P., Yang, X.L., Wang, X.G., Hu, B., Zhang, L., Zhang, W., Si, H.R., Zhu, Y., Li, B., Huang, C.L., Chen, H.D., Chen, J., Luo, Y., Guo, H., Jiang, R.D., Liu, M.Q., Chen, Y., Shen, X.R., Wang, X., Zheng, X.S., Zhao, K., Chen, Q.J., Deng, F., Liu, L.L., Yan, B., Zhan, F.X., Wang, Y.Y., Xiao, G.F., Shi, Z.L., 2020. A pneumonia outbreak associated with a new coronavirus of probable bat origin. Nature 579, 270–273.

